# Exploring morphological motifs for a single neuron based on multiple 3D reconstructions

**DOI:** 10.1101/254425

**Authors:** Jian Yang, Yishan He, Zhi Zhou, Ning Zhong, Hanchuan Peng

## Abstract

The morphology of individual neurons is useful to study structures and functions of nervous system. Researchers have invented many semi or fully automatic tracing methods to efficiently generate a reconstruction from a single neuron. Different tracing methods have different design principles, and could produce different reconstructions. However, the “common substructures” of various reconstructions, called morphological motifs, should be highly reliable. In this work, we propose a Vaa3D based framework to explore morphological motifs of 3D reconstructions from a single neuron. The framework contains four steps: (1) resampling and sorting each reconstruction according to a standard reconstruction, such as a gold standard reconstruction, consensus reconstruction or a certain reliable reconstruction; (2) applying local alignment algorithm for each pair of the standard reconstruction and a reconstruction, or each pair of two reconstructions; (3) constructing overlaps based on selected points in local alignment pairs; (4) obtaining morphological motifs by post-processing these overlaps. Under the proposed framework, three methods were implemented and tested on a dataset of 73 fruitfly neurons released by the BigNeuron project (http://bigneuron.org), which contains a gold standard reconstruction, a consensus reconstruction and about 40 automatic reconstructions for each neuron. We quantitatively evaluated these three methods to choose reliable morphological motifs.

## Introduction

The structure and function of neurons are useful for understanding the working mechanism of brains. Neuronal morphology is a major indicator for investigating neuronal structure and function, which is determined by a number of factors, including physical and biological constraints and requirements of axonal, dendritic, and network functionn^1^. In neuroscience, it is important to accurately trace, or reconstruct, a neuron’s 3D morphology. One major task of the US BRAIN Project and the European Human Brain Project is reconstruction and aggregation of neuronal morphologies on scales up to the whole rodent brain. The reconstruction of a neuron's morphology or tracing a neuron has been in practice for one century since the time of Santiago Ramóny Cajal. So far, researchers have invented many semi or fully automatic tracing methods to efficiently generate a reconstruction from a neuron. Since the morphology of neurons is so complex that any two neurons of any species have different morphologies, and the digital image of neurons is always polluted by more or less noises, it is difficult to find a tracing method which performs very well for all different types of neurons or every part of the same neuron. But for a neuron, the “common substructures” of reconstructions generated by many tracing methods with different design principles should have a high degree of reliability, and can be adopted directly. We introduce a framework to define these “common substructures” of reconstructions of a neuron, and propose three exploring methods.

In the past five decades, many computational methods and tools based on the help of computers have been developed for digital reconstruction of neurons from images^2,3,4^. Especially, two memorabilia were held to promote the research of automatic tracing methods. One is the DIADEM (short for digital reconstruction of axonal and dendritic morphology) neuron reconstruction challenge hold in 2010^5,6^. The goal of DIADEM was to develop algorithms capable of automatically converting stacks of images visualizing the tree-like shape of neuronal axons and dendrites into faithful 3D digital reconstructions^7^. More than 100 teams from worldwide registered to participate in the DIADEM Challenge. Another is a community-contributed project named BigNeuron^8^ (http://bigneuron.org). BigNeuron was launched in 2015, and its aim was to define and advance the state of the art of single-neuron reconstruction, develop a toolkit of standardized reconstruction protocols, analyze neuron morphologies, and establish a data resource for neuroscience^7,8^. The project held a series of hackathons to gather researchers and developers all over the world specifically for neuronal reconstructing and neuronal morphological studying. BigNeuron incorporated more than 30 automatic tracing algorithms, which were tested on a set of 30,000+ multi-dimensional neuronal image stacks and generated more than one million morphological reconstructions of neurons from different species (http://alleninstitute.org/bigneuron/).

So far, there are many methods or algorithms invented for tracing neuronal morphology. These methods can be categorized into semi-automatic and automatic. Semi-automatic methods require experts to delineate or validate critical parts of neuronal morphology, and other parts are reconstructed automatically by the algorithm. Most of them can generate quite accurate reconstruction results, but they are time consuming for the intervention of annotators. Automatic methods can reconstruct a neuron very quickly, even for long-range projection neurons of mouse or human. Different tracing methods design different models and have different strategies. APP (All-path pruning) is a pruning over-completed neuron-trees method^9^, which constructs an initial over-reconstruction and then simplifies the entire reconstruction by pruning the redundant structural elements. APP2 is an update version of APP^10^, and its most important idea is to prune an initial reconstruction tree using a long-segment-first hierarchical procedure. Simple Tracing uses DF-Tracing to execute a coupled distance-field (DF) algorithm on the extracted foreground neurite signal^11^. MOST ray-shooting tracing is based on the simulation of blood flow from initial seeds to compute centerlines and their corresponding radii^12^. SmartTracing invokes a user-provided existing neuron tracing method to produce an initial neuron reconstruction, and then uses a machine learning framework to predict neuron signal in all image area^13^. Rivulet is based on the multi-stencils fast-marching and iterative back-tracking, and can trace discontinuous areas without being interrupted by densely distributed noises^14^. There are many other 3D automatic tracing algorithms, such as Rayburst sampling based approach^15^, automatic contour extraction method^16^, Open-Curve Snake^17^, probabilistic approach with global optimization^18^, ray casting^19^, tube-fitting model^20^, tubularity flow field (TuFF) method^21^, 3D tubular model^22^, SmartScope2^23^, M-AMST^24^ and so on.

However, automatic tracing methods developed for different application scenarios and based on different models and strategies typically have varying performance, especially while being used on neuron images of variable quality and different species^8^. Because most of these methods have not been directly cross-tested thoroughly, it is unclear which methods are best matched with different imaging modalities or datasets^7^. Even if there exists a best automatic tracing method, it is hard to guarantee its reconstruction for a neuron is good everywhere. One method might perform well at some parts of the image and another method might be good at other parts. For the image of a neuron, reconstructions generated by different methods always have some similar parts and dissimilar parts. These similar parts reflect a high degree of agreement that many methods with different models or strategies reach. Therefore, it is reasonable to suppose that these similar parts are good substructures for tracing the neuron’s morphology. We call these similar parts morphological motifs of reconstructions of different methods. Morphological motifs can be used to generate a gold standard reconstruction of a neuron. In practice, in order to obtain morphological reconstruction with high-accuracy, neuronal reconstructions are often still made or checked segment by segment manually by human experts with the help of computers. Morphological motifs can be omitted in the manual tracing or checking course.

Actually, common substructures in morphologies of different neurons are called motifs in computational biology. Wan et al. developed BlastNeuron to compare neurons in terms of their global appearance, detailed arborization patterns, and topological similarity^25^. The local alignment of BlastNeuron is able to find the corresponding branches or sub-structure of neuron morphologies for a pair of the tightly connected neurons, and pinpoints structure motifs of two similar neurons. But morphological motifs of much more than two reconstructions of a neuron need to be redefined and studied further. In addition, there is the definition of topological motifs of reconstructions of different neurons^1,26^. Gillette and Ascoli^1^ decomposed the neuron topology into sequences of branching patterns, and then proposed a method to compare neuron structures using sequence alignment. The method is able to identify the difference in branching patterns in dendritic and axonic arbors and extracting common topological “motifs” in the structure^1,25^. Topological motifs use only topology of neurons and its aim is to find common topological structure in different neurons, but morphological motifs is based on geometry of reconstructions and its goal is exploring common geometric morphology hidden in various reconstructions.

## Method and Results

### Overview of the framework

For a neuron, many different reconstructions can be conveniently obtained by implementing various automatic tracing methods as plugins in Vaa3D. Similar parts of these reconstructions are good parts for tracing the neuron’s morphology, and are called morphological motifs. Morphological motifs contain both detailed structural and spatial characteristics, and can represent certain substructures of a single neuron. In the framework, all data of a single neuron includes its light-microscopic image, a gold standard reconstruction, a consensus reconstruction and about 40 various automatically traced reconstructions. All these reconstructions are presented by SWC format^27^, in which neuron morphologies is described as tree structures with location, node’s radius, parent node and some other attributes. The overview of the framework and one of its results for reconstructions of a single neuron is demonstrated in Fig. 1. Taking a standard reconstruction (it may be a gold standard reconstruction, a consensus reconstruction or just a good tracing reconstruction) and all reconstructions generated by various tracing algorithms as input, our framework does some pre-processing for standardization at first, and constructs local alignment pairs between any two of reconstructions by the local alignment algorithm in BlastNeuron^25^ (step S1 in Fig. 1). Then, local alignment pairs are pruned (step S2 in Fig. 1) and overlapping points are obtained (step S3 in Fig. 1). Finally, they are connected to a tree-like morphological structure and post-processed to give morphology motifs (step S4 in Fig. 1).

**Figure 1.**
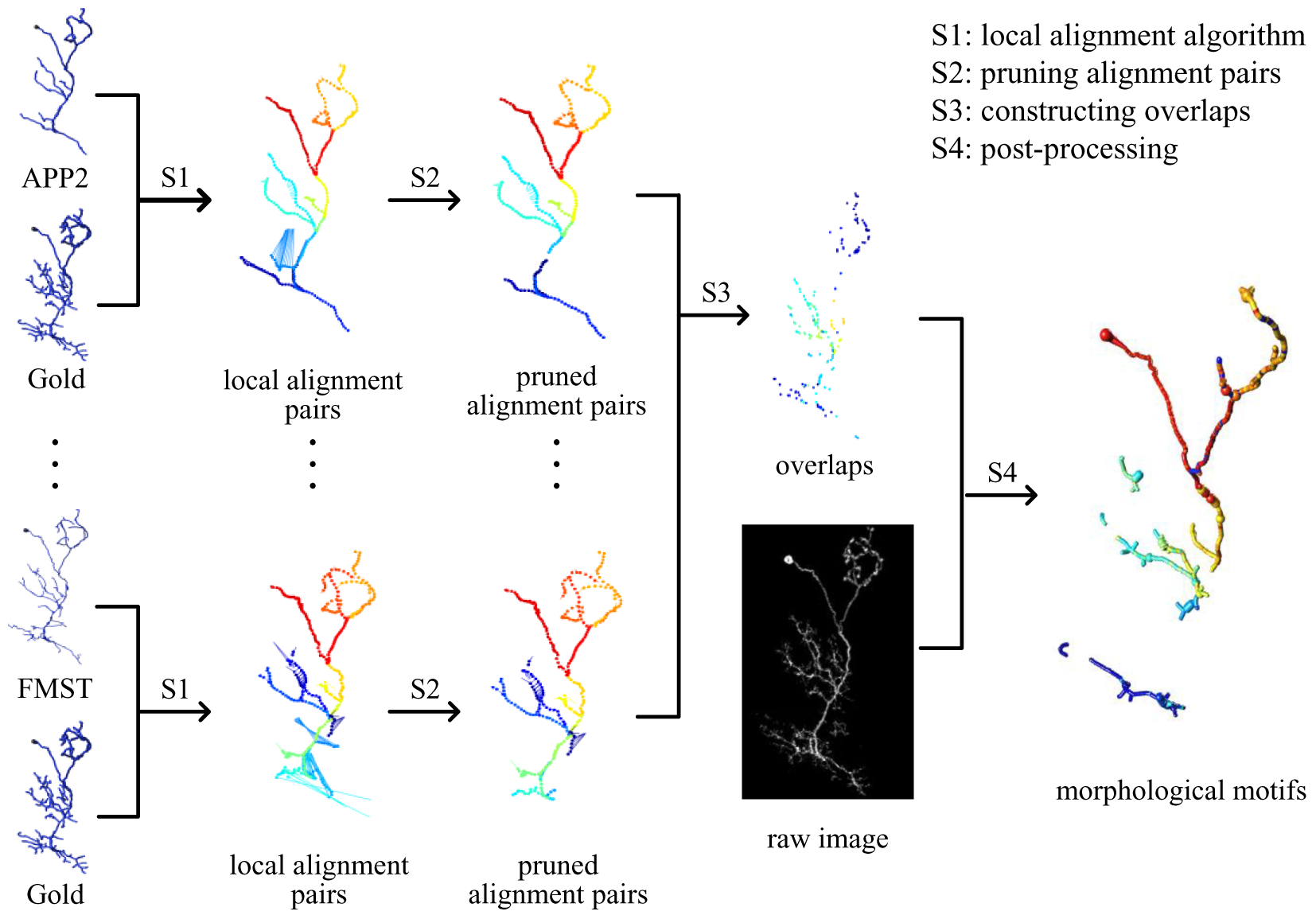
Overview of the proposed framework. Local alignment pairs between the gold reconstruction and an automatic reconstruction are found (S1) and pruned (S2), and overlaps of pruned local alignment pairs corresponding to all automatic reconstructions are constructed (S3). After some post-processing steps (S4), morphological motifs of all reconstructions are obtained.

### Pre-processing reconstructions

Neuron tracing algorithms use different methods and generate quite different reconstructions, which might be good or rather bad, especially for complex neuron morphological structures of different kinds of neuron dataset. It is necessary to implement some pre-processing steps for standardizing all reconstructions and remove some unreasonable results. Pre-processing steps contain resampling, sorting and filtering, which can be carried out by running corresponding plugins of Vaa3D^28,29^.

Reconstructions of neuron tracing algorithms always have different node density on the neuron tree. In order to limit its impact, we resample all reconstructions with a fixed step length by running resample_swc function (https://github.com/Vaa3D/vaa3d_tools/tree/master/released_plugins/v3d_plugins/resample_swc). Then node numbers of each resampled reconstruction can truly reflect the size of its neuronal tree. For satisfying the requirement of the local alignment algorithm, we use sort function (https://github.com/Vaa3D/vaa3d_tools/tree/master/released_plugins/v3d_plugins/sort_neuron_swc) to reconnect all reconstructions and turn them into a single tree if some of them have multiple trees. Meanwhile, we reset the root node of a reconstruction to its node with the minimum distance to the root node of a standard reconstruction (gold reconstruction, consensus reconstruction or APP2 reconstruction). At last, we filter out some under-fitting and over-fitting reconstructions with too few or too many nodes compared to their average value. Especially, over-fitting reconstructions have a great impact on the subsequent steps and the final morphological motifs. The number of nodes in remained reconstructions satisfies the following boundary condition:

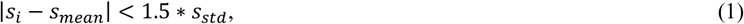

where *s*_*i*_ is the node number of the *i*th reconstruction, *s*_*mean*_ and *s*_*std*_ are the aveerage and standard deviation of node numbers of all reconstructions.

### Generating local alignment pairs

The local alignment algorithm which compares neurons’ morphology locally at the compartment level is the most important component of BlastNeuron^25^. It finds the corresponding relationship of segments in two reconstructions by matching their topology and geometry and plays a key role in our framework.

The inputs of the local alignment algorithm are two neuron reconstructions (denoted by A and B, respectively) having tree-topology structures. Both reconstructions are firstly normalized to the same center location in 3D space by using resampling and PCA (Principal Component Analysis) method, and then the algorithm constructs a Euclidean distance matrix of each point of A to all other points of B. Next, it uses a RANSAC sampling process^30^ to estimate the optimal affine transform from A to B, so the correspondence of A and B can be found in a same space.

Considering the shape and location of neuronal arbors, the algorithm decomposes reconstructions into several simple line segments bounded by branching nodes and tip nodes. Through calculating the distance of these segments and their inner nodes using dynamic programming, the corresponding nodes are matched and then local alignment pairs are formed.

### Consensus of reconstructions

Consensus reconstructions are constructed by the BigNeuron community automatic (https://github.com/Vaa3D/vaa3d_tools/tree/b9348967b08a27029929df50d9c8f929d07340f4/hackathon/xiaoxiaol/consensus_skeleton_2). The consensus is extracted from several reconstructions of a neuron and involves almost all morphological structures in each tracing results. For a neuron, if there is no gold standard reconstruction, consensus reconstruction could be an appropriate candidate as the standard reconstruction in our framework.

The basic idea of generating a consensus reconstruction is merging multiple neuron reconstructions via clustering. To speed up the calculation of the clustering process, each candidate reconstruction is represented by kd-trees first. For each node in each candidate reconstruction, its nearest nodes in all other candidates are found, and the node is moved to the center of those nearest nodes. That is to say, each candidate reconstruction is moved to the center of other candidate reconstructions. The next step is merging these centralized candidates to a list of consensus nodes with vote information. Finally, consensus nodes are connected using edges votes (collected from original individual neuron trees) via minimum spanning tree algorithm. Meanwhile, some unconfident branches are trimmed and some short branches with little significance but relative high effect on the final result should be paid more attention.

### Constructing overlaps with three methods

If we have a given standard reconstruction (a gold reconstruction manually generated by human experts or a consensus reconstruction based on all automatic reconstructions), local alignment pairs could be found between it and other reconstructions. If we have no standard reconstruction, local alignment pairs are found between each pair of reconstructions. We denote the former case by “standard reconstruction” and the latter case by “all reconstructions”. In “standard reconstruction”, the standard reconstruction will be used one time for each of other reconstruction. While defining the overlapping areas of local alignment pairs, overlaps might be defined on the standard reconstruction or center points of line segments connecting local alignment pairs. We denote these two cases by “reference manner” and “mean manner”. Since “all reconstructions” case could not match to “reference manner”, we have total three cases according to different combination of “standard reconstruction”, “all reconstructions”, “reference manner” and “mean manner”. As showed in Table 1, three methods corresponding to these three cases are abbreviated to SR, SM and AM, respectively.

**Table 1.** Combination manners and names of the proposed three methods.

|  | reference manner | mean manner |
| --- | --- | --- |
| standard reconstruction | SR | SM |
| all reconstructions | -- | AM |

Matched local alignment pairs found by BlastNeuron consist of line segments between two nodes on corresponding to two reconstructions, the shorter the better matching. We discard these pairs with length larger than a given threshold and discuss two defining manners of overlaps only on remained short pairs.

For SR method, a given standard reconstruction is used to find local alignment pairs and each matched local alignment pair has a node on the standard reconstruction. That node is used to represent the corresponding local alignment pair. As shown in Fig. 2 (a), standard, A and B are the standard reconstruction and other two reconstructions. A red-filled circle represents a local alignment pair. The distance between two local alignment pairs is calculated by the Euclidean distance between their representative nodes on the standard reconstruction (red-filled circles). For two local alignment pairs corresponding to two reconstructions, if their distance is less than a given threshold (called neighbor distance threshold in experiments), their representative nodes are put into the overlapping set of these two reconstructions. For a given threshold number *n* (called count number threshold in experiments), if a representative point belongs to the overlaps of at least *n* pairs of reconstructions, the node is put in the final overlaps of all reconstructions.

**Figure 2.**
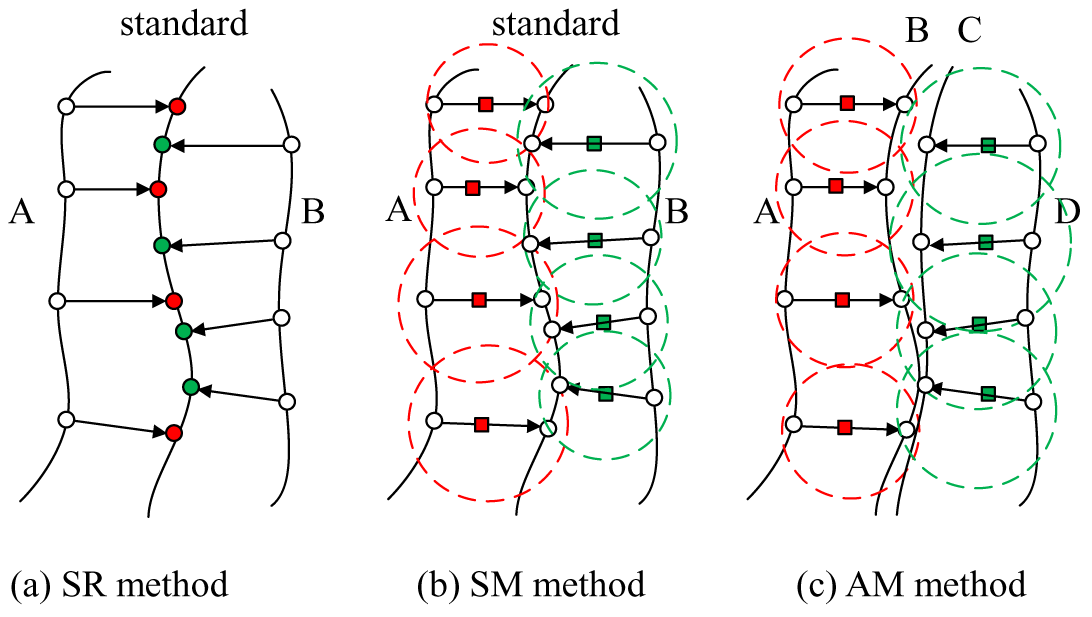
Three methods for constructing overlapping points set. “standard” is a standard reconstruction, and A, B, C and D are different reconstructions. Unfilled circles are nodes on reconstructions, arrows represent local alignments between two nodes, and filled circles and filled squares (red and green) are representative points of their corresponding local alignment pairs. These filled circles and squares are also candidate overlapping points in SR, SM and AM method. For dotted circles (red and green) in (b) and (c), their centers are filled squares and their radii are the centers’ radii estimated from the original light-microscopic image.

For SM method, the center point of the line segment of a local alignment pair is taken as its representative point. As shown in Fig. 2 (b), green-filled squares represent local alignment pairs of between a reconstruction and standard, respectively. Then we estimate the radius of each representative point in the original light-microscopic image of the neuron. In Fig.2 (b), centers of dotted circles locate at representative points and their radii are radii of representative points. The distance between two local alignment pairs is also calculated by the Euclidean distance between their representative points (red solid circles and green solid squares). For two local alignment pairs between two reconstructions, if their distance is less than the sum of the radii of their representative points, their representative points are put into the overlapping set of these two reconstructions. Then with a given count number threshold, the final overlaps of all reconstructions can be defined as SR method.

For AM method, there is no standard reconstruction, and representative points of local alignment pairs can be found as SM method, as shown in Fig. 2 (c). Based on these representative points, the overlapping points of two pairs of reconstructions in AM method and the final overlaps of all reconstructions are defined as SR method.

### Post-processing overlaps

All points in overlaps constructed by SR method are selected from nodes of the standard reconstruction, so they can be processed efficiently. But for SM and AM methods, overlaps points are selected from center points of line segments of local alignment pairs. There might be too many near points in the overlaps, so we estimate these points’ radii and trim those points covered by other points in the overlaps. Post-processed overlapping points compose the skeleton of the final morphological structure. We reconnect them into tree-like structures and estimate the nodes’ radii using the light-microscopic image of a single neuron. To generate meaningful morphological motifs, we sort these select points to produce some trees, estimate each node's radius on these trees and remove short trees (with no more 3 nodes).

### Visual demonstration of generated morphological motifs

The proposed framework with its three methods was tested on reconstructions of 73 fruitfly neurons. The dataset was released by the BigNeuron community. Each neuron has a gold reconstruction produced manually by human experts, a consensus reconstruction and other about 40 reconstructions generated by automatic tracing methods with different parameters. The first line of Fig. 3 demonstrates some reconstructions of a neuron including gold standard reconstruction (Gold), consensus reconstruction (Consensus), and reconstructions of APP2, Neutube, EnsembleNeuronTracerV2n, SmartTracing and FMST. The gold standard reconstruction and the consensus reconstruction have very similar skeletons, and they are only different at some short branches. The consensus reconstruction is quite good. Other reconstructions also reflect the skeleton of the neuron, but some of them miss some important branches due to excessive pruning and some of them have redundant branches caused by lacking trimming.

**Figure 3.**
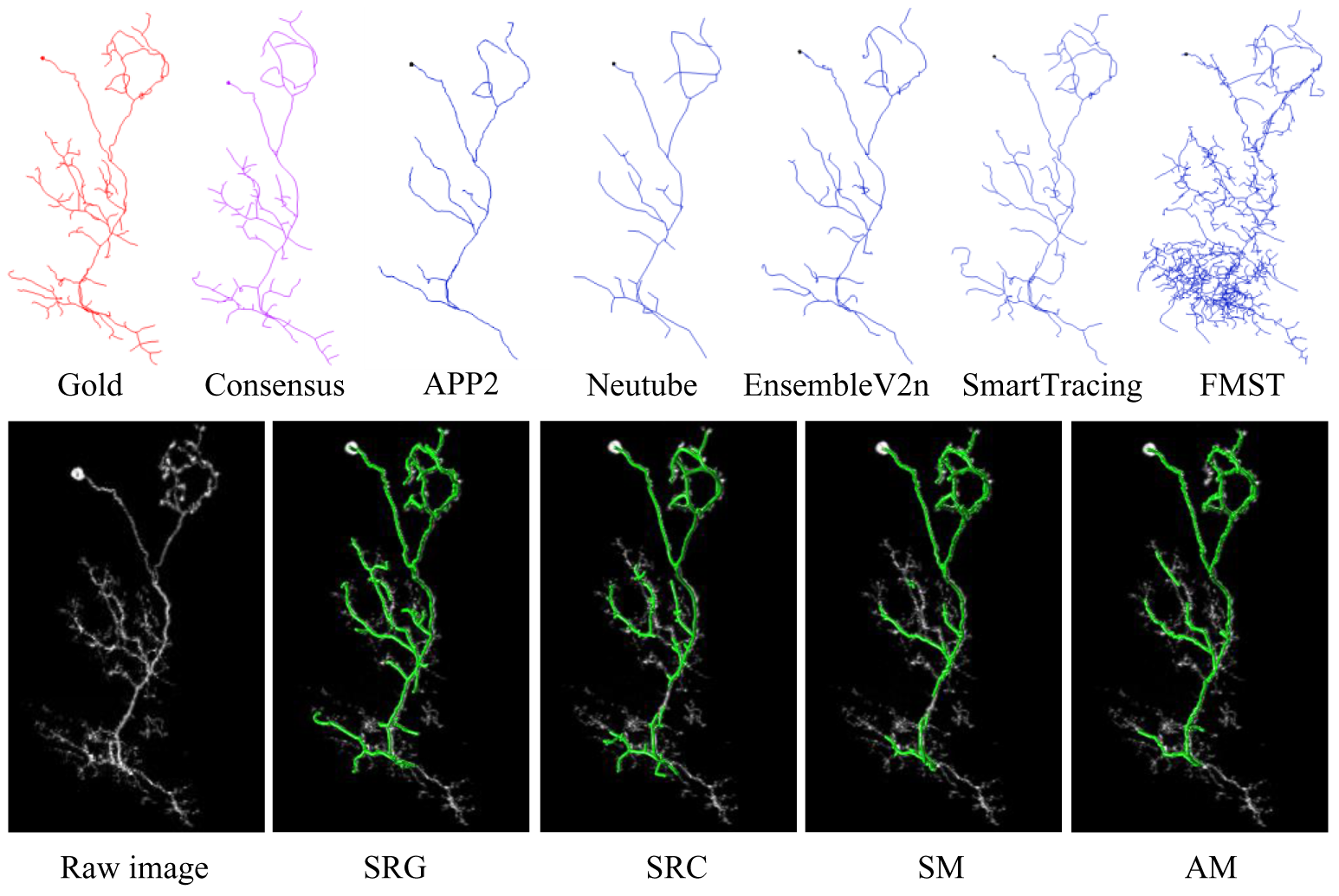
Morphological motifs of reconstructions for a fruitfly neuron. Subfigures in the first line are gold reconstruction, consensus reconstruction and 5 of 40 reconstructions generated by automatic methods. Raw image of the neuron and morphological motifs of these 40 reconstructions generated with SR method (SRG denotes taking its gold standard reconstruction and SRC denotes taking consensus reconstruction as the standard reconstruction), SM method and AM method are demonstrated in subfigures in the second line, where green lines are morphological motifs.

For each neuron, its gold standard reconstruction and consensus reconstruction were taken as the standard reconstruction in SR method, and denoted by SRG and SRC, respectively. Its consensus reconstruction was taken as the standard reconstruction in SM method, and its APP2 reconstruction was used to pre-process other reconstructions in AM method. So with a given group of parameters in each method, we obtained four morphological motifs of these reconstructions by implementing the proposed framework. These four results for a neuron and its raw image were illustrated in the second line of Fig. 3. In these subfigures, image background is the gray image of a neuron, and green lines are morphological motifs corresponding to different methods. These morphological motifs are “common” parts of many reconstructions, and have high reliability.

### Morphological motifs generated by different methods with different parameters

The framework with different methods or with same methods but different parameters generated different morphological motifs for a same group of reconstructions. To investigate these differences, we implemented the framework with each method (or each standard reconstruction) under several different groups of parameters. Six groups of parameters for SR method and three groups of parameters for SM method and AM method were implemented. For SR method, the neighbor distance threshold was set to 3 and 5, and the count number threshold was set to 4, 6 and 8. Morphological motifs denoted by SRG_3_4 and SRC_3_4 are results of SR method while taking the gold standard reconstruction and consensus as the standard reconstruction respectively, 3 as the neighbor distance threshold and 4 as the count number threshold. For SM method and AM method, the count number threshold was set to 12, 15 and 18. The left three columns of Fig. 4 illustrate morphological motifs generated by three methods and three groups of parameters for a fruitfly neuron. It can be seen that for a method, the larger the neighbor distance threshold is or the smaller the count number threshold is, the more nodes morphological motifs have.

**Figure 4.**
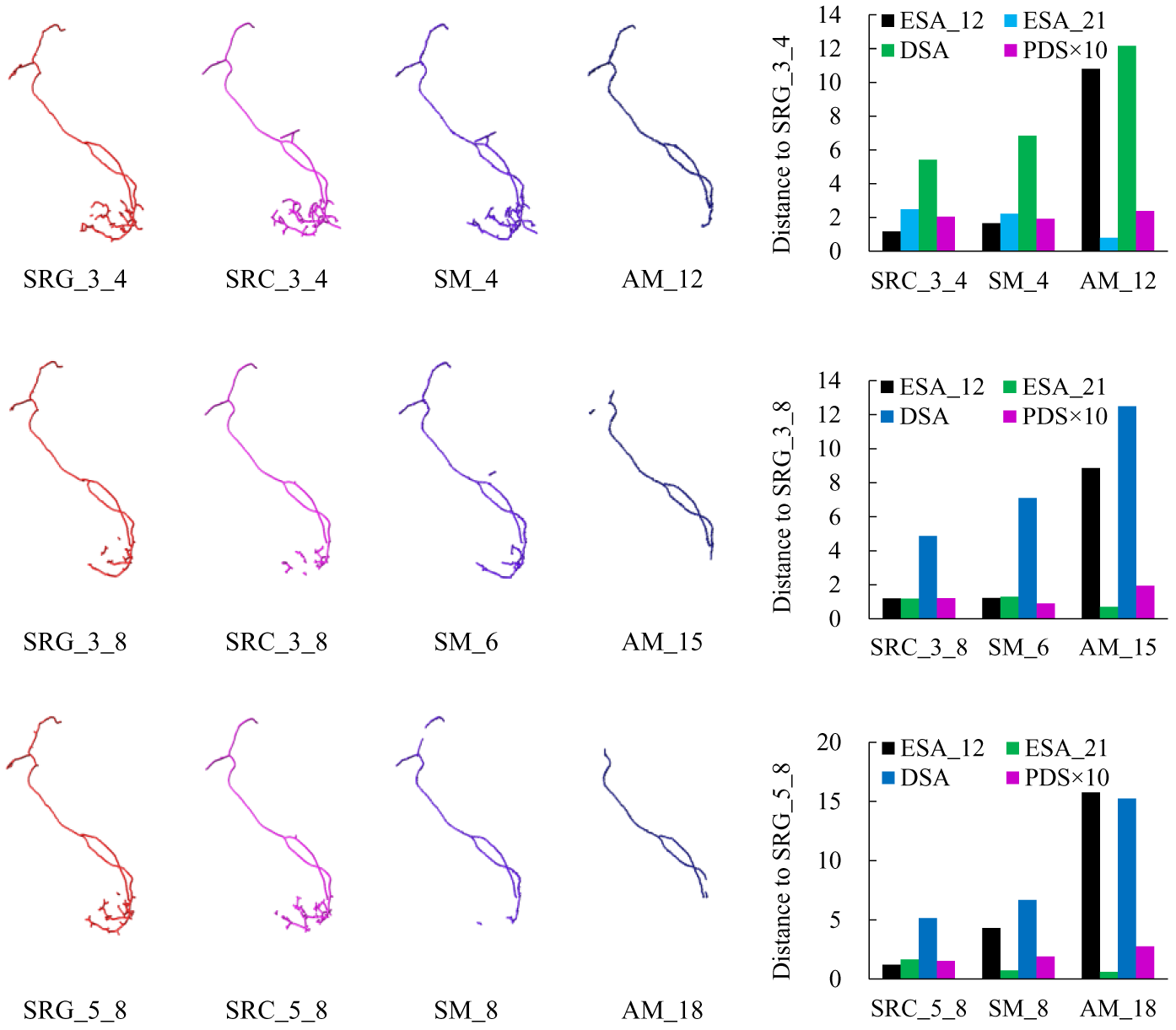
Morphological motifs generated by different methods with different parameters for a fruitfly neuron. Subfigures in column 1 to 4 are morphological motifs generated by SR method with the gold standard reconstruction as the standard reconstruction, SR method with the consensus reconstruction as the standard reconstruction, SM method and AM method, respectively. For each method (or each standard reconstruction), three results with different parameters are demonstrated in row 1 to 3. The last column of each row is the bar graph of four distances (ESA_12: entire-structure-average from neuron 1 to 2, ESA_21: entire-structure-average from neuron 2 to 1, DSA: different-structure-average and PDS: percent of different-structure ESA_12) between the SRG result and other three methods’ results on the same row, where values of PDS (between 0 and 1) are enlarged ten times for a better show.

To quantitatively investigate differences between motifs generated by different methods, four distance measures were calculated for morphological motifs in Fig. 4 and motifs of 73 neurons’ reconstructions. They are entire-structure-average from neuron 1 to 2 (ESA_12), entire-structure-average from neuron 2 to 1 (ESA_21), different-structure-average (DSA) and percent of different-structure (PDS)^31^, and are functions of a neuron distance plugin in Vaa3D. The bar graph of these distances between the SRG based motifs and other three motifs in each row of Fig. 3 were plotted in the last column of Fig.3. We calculated these distances between gold standard reconstructions based morphological motifs and other motifs, and their means on 73 fruitfly neurons were plotted in Fig. 5, where gold denotes results of SR method while taking the gold standard reconstruction as the standard reconstruction and SRC denotes results of SR method while taking consensus reconstruction as the standard reconstruction. The smaller these scores are, the more similar gold standard reconstructions based motifs and other motifs are. From Fig. 5, we can see that consensus reconstructions based motifs SRC_3_4 and SRC_5_4 are relatively more similar to gold reconstructions based motifs. That is to say, for this dataset, if we do not have gold reconstructions, we may use results of SRC_3_4 and SRC_5_4 to substitute its motifs. But for other datasets, which method and which parameters we should select while implementing the framework still need more investigations.

**Figure 5.**
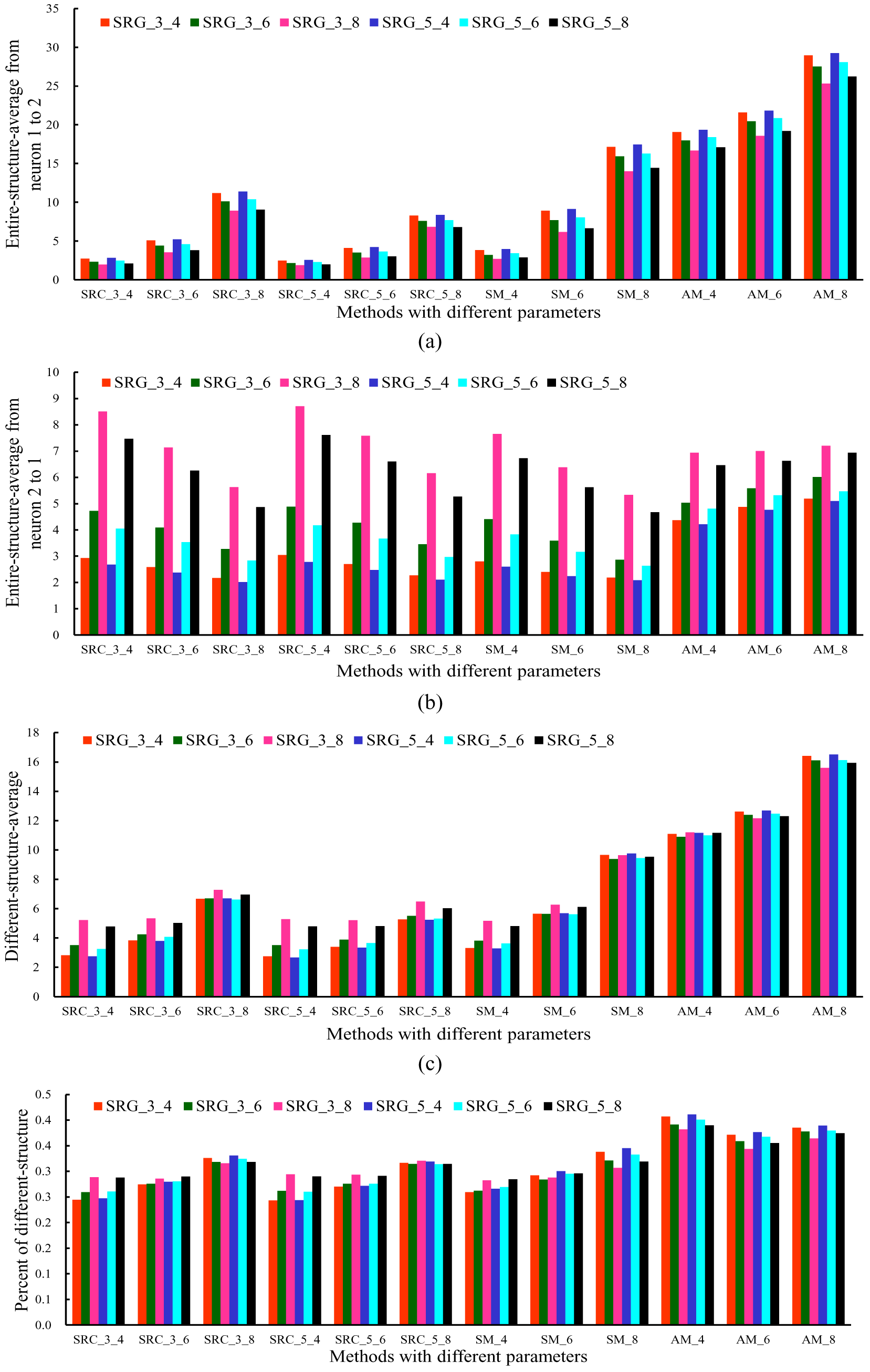
Means of four distances between gold standard reconstructions based morphological motifs and other motifs. Subfigures in (a)-(d) are means of entire-structure-average from neuron 1 to 2, entire-structure-average from neuron 2 to 1, different-structure-average and percent of different-structure, respectively.

## Discussions

We developed a software pipeline to implement three different methods for finding morphological motifs of reconstructions. Morphological motifs are similar and common substructures among multiple reconstructions, and are highly reliable parts of multiple automatic tracing approaches with different models or methods. For a large neuron image, we obtained many reconstructions by implementing all kinds of automatic tracing approaches, and then found their morphological motifs using our framework and methods. These motifs can be used without manual checking while constructing a gold standard reconstruction for the neuron.

The overall motifs might consist of several broken segments with unequal length, and motifs produced by different methods are different. SR method based motifs depend more on the given standard reconstruction and are always bigger than motifs of SM method and AM method. AM method is independent of standard reconstruction and only utilizes all kinds of automatic tracing results. It uses least intervention of human factors in our framework, but its running time is too long. For *n* reconstructions, we calculated *n*×(*n-*1) pairs of local alignment pairs, and only selected about 3% (20) pairs to construct their overlaps. A better manner for reducing its time complexity is to be considered.

While visually checking morphological motifs for these 73 fruitfly neurons, we find that all their motifs coincide with their skeleton or morphology structure and mainly locate at simple neuronal segments, which are relatively easy to reconstruct for most tracing methods. Especially for SM method and AM method, the final motifs almost only locate at the trunk of neuronal trees and have few branches. Image regions containing multiple branches are difficult to tracing their structures and different tracing methods generate quite different results. It is impossible to count on that a tracing method can perform well everywhere in these complex regions, so tracing results at these regions should be checked carefully or reconstructed by human experts.

Some neurons of same category have similar morphology, and morphological motifs of reconstructions for different neurons might contain some special morphology features of that kind of neurons. Though location difference of reconstructions for multiple neurons may be settled by translation, rotation and other pre-processing, morphological motifs of multiple neurons might be very small. Local structures of multiple neurons are quite different and local alignment pairs are too strict to evaluate. This results in that our framework has not enough local alignment pairs as its input. Some more sophisticated characterization of local similar structure might be the solution of the issue, but it needs to be studied further.

## Acknowledgements

The authors thank the BigNeuron community for providing the experimental data. This work is partially supported by the National Basic Research Program of China (No. 2014CB744600) and the National Natural Science Foundation of China (No. 61420106005).

## Author contributions

Hanchuan Peng designed the main idea of the project, and Jian Yang, Zhi Zhou and Ning Zhong took part in its discussion. Yishan He and Zhi Zhou programed the proposed methods. Jian Yang and Yishan He prepared the figures with the help of others. Jian Yang and Yishan He wrote the main manuscript text. All authors revised the manuscript.

## Data availability

The datasets generated during and/or analyzed during the current study are available from the corresponding author on reasonable request. The source code of our framework and its three methods is openly distributed along with Vaa3D code repository (https://github.com/Vaa3D/vaa3d_tools/tree/master/hackathon/heyishan/blastneuron_bjut).

